# The claustrum drives large-scale interactions of cortical circuits relevant to long-term memory

**DOI:** 10.1101/2023.03.02.530783

**Authors:** S. Mutel, JR. Renfer, I. Rodriguez, A. Carleton, RF. Salazar

**Author notes:** Equal contribution as first authors (S. Mutel, JR. Renfer). Equal contribution as last authors (I. Rodriguez, A. Carleton, RF. Salazar).

## Abstract

The consolidation and recall of episodic memories rely on distributed cortical activity. The claustrum, a subcortical structure reciprocally connected to most of the cortex, may facilitate inter-areal communication necessary for these processes. We report here that the functional inhibition of claustral projection neurons affects directional interactions and the coordination of oscillatory neuronal patterns in the fronto-parietal network. Moreover, the inhibition of these neurons has a detrimental effect on concurrent oscillatory events relevant to the consolidation of contextual fear memory. Last, we demonstrate that biasing the directional flow of information between the latter two cortical areas enhances the retrieval of a remote contextual memory. We propose that the claustrum orchestrates inter-areal cortical interactions relevant to contextual memory processes by affecting the latency of neuronal responses.

**One-Sentence Summary:** The claustrum coordinates inter-areal cortical activity.

## Introduction

Our ability to remember numerous events during our lifetime remains a mysterious property of the brain. Recently, exciting new findings suggest a storage space allocated in a specific layer of the cortex (*1, 2*). These findings are consistent with the fact that memory processes are distributed throughout the whole cortex (*3*). However, episodic memories change over time and these changes rely on neuronal interactions (*4, 5*). Local neuronal interactions relevant to learning-related synaptic plasticity have been studied (*6, 7*). In contrast, we still don’t know how distant cortical interactions, potentially binding pieces of memories supported by opposite locations in the cortex, may occur. One structure, the claustrum, is reciprocally connected with most of the cortex (*8, 9*), with a bias toward associative areas (*10*). In rodents, identifying genetic markers enabled the specific targeting of claustrum neurons (*10–12*) and the demonstration of a role played by the latter in attentional processes (*13–15*), learning (*16*) and sleep (*17, 18*). However, little is known about the role of the claustrum in interregional cortical interactions related to memory processes.

### The claustrum regulates fronto-parietal communication

To assess the potential role of the claustrum in regulating inter-areal cortical interactions, we chemogenetically reduced the excitability of claustral glutamatergic projection neurons while monitoring cortical activity (see Fig. S1A for examples of electrode locations). To this aim, we used *Vglut2*-cre mice, which strongly express the recombinase in projection neurons of the claustrum (*19*). The inhibitory designer receptor exclusively activated by designer drugs (hM4D DREADD) was virally expressed in *Vglut2*-positive claustral neurons of transgenic mice (Fig. 1A), and electrophysiological signals in the prefrontal (PFC) and retrosplenial cortex (RSC) were analyzed during periods of quietness (Fig. 1B). Injections of clozapine N-oxide (CNO; also performed in control mice) reduce the excitability of neurons expressing hM4D DREADD, a.k.a. inhibition, but not in neurons expressing the control compound lacking the DREADD receptor (mCherry) (*20*). Inhibiting *Vglut2*-positive claustral neurons did not alter oscillations power in the PFC and RSC (Fig. S1B) but affected the flow of cortical information. Indeed, directional interactions between PFC and RSC, assessed using the Wiener-Granger causality (*21–23*) on pairs of local field potentials (LFP; Fig. 1B inset), were strongly diminished by an inhibition of *Vglut2*-positive claustral neurons (Fig. 1C and Fig. S1). This effect was more accentuated in signals from the PFC to the RSC than the inverse relationship (Fig. 1C, D). In the gamma frequency range (30 to 120 Hz) and at High-Frequency Oscillations (HFO; 120 to 250 Hz), signals from the PFC to the RSC tended to be dominant compared to the opposite direction but the inhibition of *Vglut2*-positive claustral neurons reverse the dominance pattern (Fig. 1D). Interestingly, in a few recordings from the claustro-insular region, *Vglut2*-positive claustral inhibition tended to reduce the signals from the claustro-insular region to the RSC both in the gamma and HFO frequency bands (Fig. S2). Thus, *Vglut2*-positive claustral neurons modulate inter-areal communication in the fronto-parietal network.

**Figure 1.**
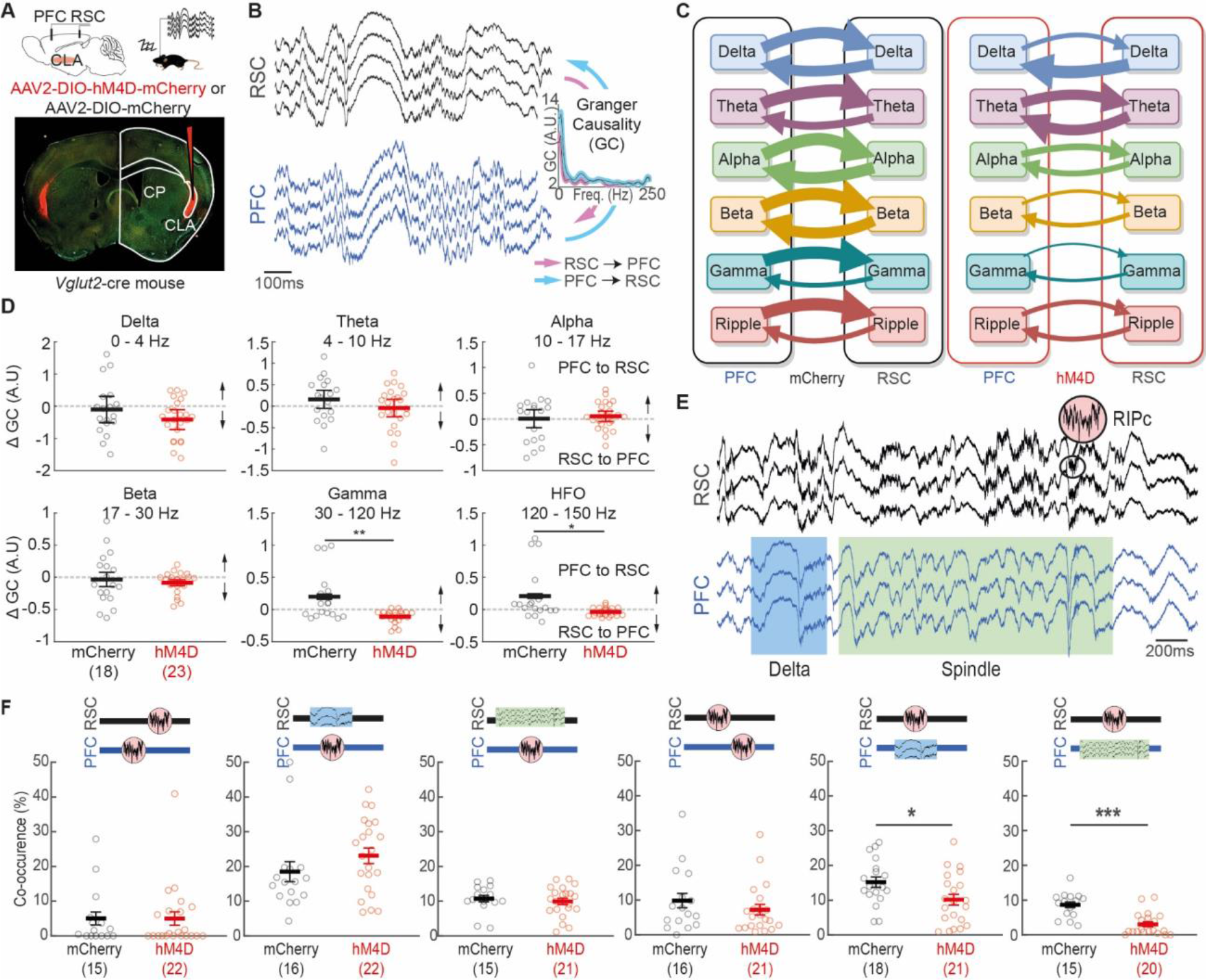
The claustrum influence fronto-parietal communication. (**A**) Claustral *Vglut2*-cre neurons were infected with AAV vectors conditionally expressing either hM4D-mCherry DREADD or mCherry alone. Tetrodes were placed in the PFC and RSC. (**B**) Example traces of simultaneously recorded signals. Spectral Wiener-Granger causality (GC) measured between the two cortical areas. (**C**) Claustral inhibition tampers down fronto-parietal directional interactions. The width of the arrows represents the mean of the population (n = 23 and 18 for hM4D and mCherry treatments, collected in 11 and 9 mice, respectively). (**D**) At high frequency bands, the inhibition of *Vglut2*-positive claustral neurons biased fronto-parietal directional interactions. (**E**) Example traces of simultaneous signals. Representative examples are shown for ripples (RIPc; see pink inset) in RSC, a delta wave (blue rectangle), and a spindle (green rectangle) in PFC. (**F**) Interareal co-occurrence of RIPc events, delta waves and spindles are presented separately for hM4D or mCherry-treated mice. In all panels, significance in the graphs was marked for p-values < 0.05 (*) and 0.001 (***). In all Figures, the number of observations n is displayed in parenthesis, statistical details are presented in the Supplementary Text and error bars represent the mean and the SEM.

High-frequency signals in the hippocampus nest oscillatory events known as sharp-wave ripples. These events reflect specific brain-wide network activity (*24–26*). The coordination between different cortical areas and the hippocampus operates via complex activations of various oscillatory events such as ripple-like high-frequency oscillations (RIPc, 100-150 Hz), delta waves (0-6 Hz), and spindles (9-17 Hz) (*27–30*). These events occur in the RSC and in the PFC and are related to hippocampal sharp-wave ripples (*24, 28, 31-33*). Thus, we explored whether the claustrum coordinates such oscillatory events in the fronto-parietal network. We first detected delta waves (as specific events, not to be confound with the delta frequency band), RIPc and spindles in recordings from the PFC and the RSC (Fig. 1E). Claustrum chemogenetic inhibition had little impact on the duration and frequency of these events, except for delta waves being increased in the PFC (Fig. S3A, B). In contrast, claustral manipulation reduced drastically the occurrence of RIPc in the RSC during ongoing delta waves or spindles in the PFC (Fig. 1F). Interestingly, this effect was unilateral as the incidence of RIPc in the PFC, either during a delta wave or during a spindle in the RSC, was unaffected by the same manipulation. Altogether, these data indicate that the unilateral coordination of RIPc, delta waves, and spindles within the fronto-parietal network strongly depends on claustrum integrity.

### Contextual fear memories rely on claustrum-dependent fronto-parietal coordination

To evaluate the functional consequence of claustrum perturbations, mice were submitted to classical Pavlovian fear conditioning. *Vglut2*-positive claustral inhibition was selectively induced either during the conditioning using an optogenetic approach (Fig. 2A-C) or during the consolidation period after the conditioning using a chemogenetic procedure (Fig. 2D-F). Both manipulations reduced the freezing response when mice were exposed to the conditioning context in the following days. The reduction was accentuated over the course of a month but did not occur either when a saline solution was injected instead of CNO (Fig. 2E) or when the electric foot-shock was not paired with light stimulation (Fig. S4A). Furthermore, no deficiency in freezing behavior or associative abilities was induced by claustral manipulations, as the freezing response to the conditioned tone was unaffected (Fig. 2C and F). One potential explanation for the contextual memory deficits may be an influence on behavioral activity and its competition with memory processes. However, it is unlikely as CNO activation of hM4D DREADD did not affect locomotion in an open field area (Fig. S4B) or clean cages (Fig. S4C). On the contrary, we even observed that claustral inhibition favored the total duration of stillness in home cages (Fig. S4D). To further characterize claustrum-related deficits in contextual memory retrievals, we investigated whether the functional alteration of specific claustro-prefrontal or claustro-retrosplenial projections could phenocopy such deficits. Using a retrograde viral cre delivery combined with local injections of cre-dependent hM4D DREADD in wild-type animals (*34*), the latter projections were inhibited during the memory consolidation period. None of these manipulations altered memory consolidation processes (Fig. S5). Thus, reducing the excitability of *Vglut2*-positive claustral projection neurons either during fear conditioning or during its memory consolidation period disrupts long-term contextual memory.

**Figure 2.**
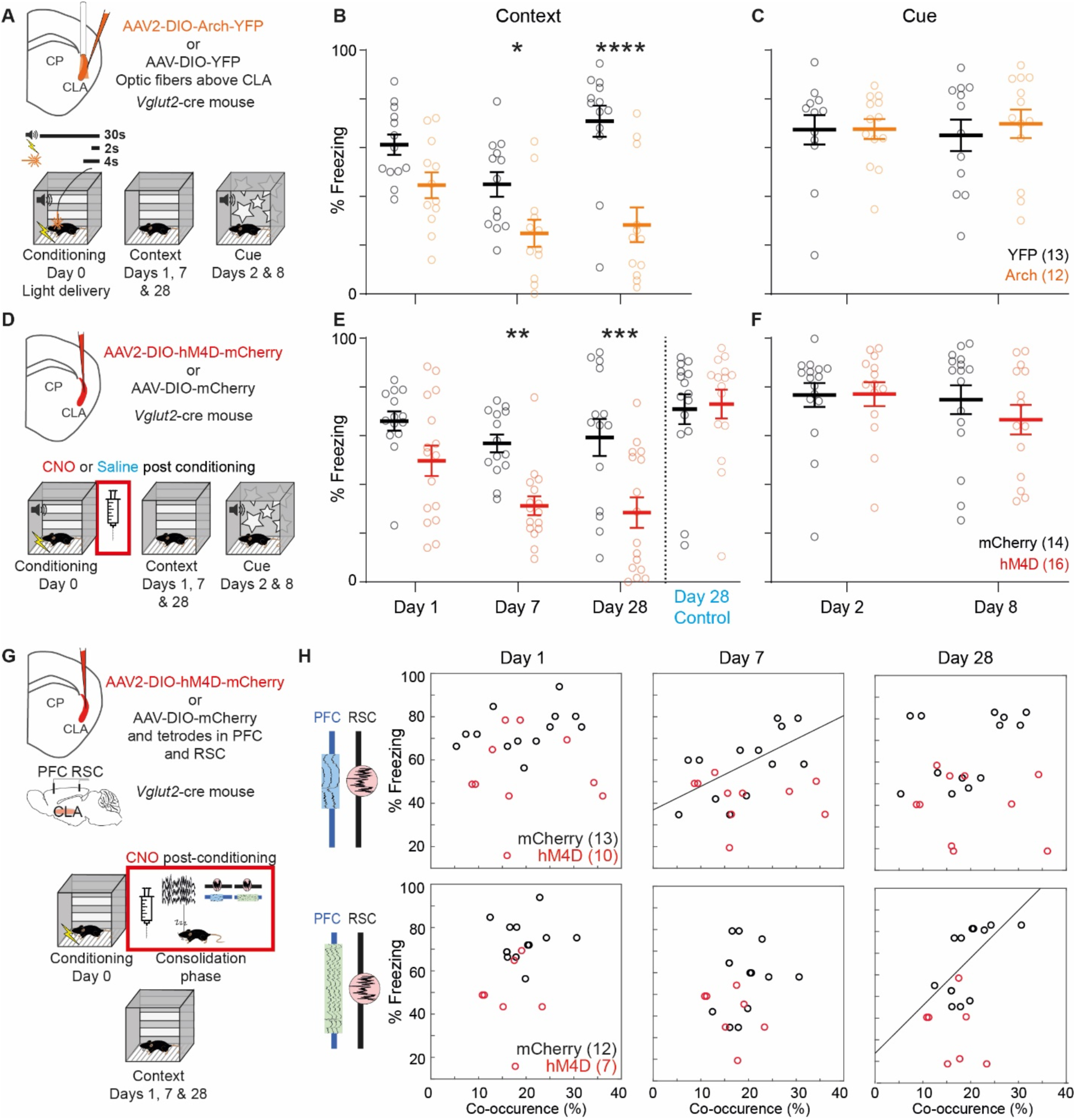
Claustrum-mediated alteration of fronto-parietal communication during consolidation periods impairs long-term memory. (**A**) The CLA of *Vglut2*-cre mice was infected with a AAV virus containing either an inhibitory opsin (Arch) or not (YFP). (**B-C**) Freezing response to the CC (CS) on days 1 (2), 7 (8), and 28 post-conditioning. (**D**) The claustrum of *Vglut2*-cre mice was infected with AAV vectors conditionally expressing either hM4D-mCherry DREADD or not. In an additional experiment, a 0.9% saline solution was injected instead of CNO (blue text). (**E-F**) Freezing response to the CC (CS) on days 1 (2), 7 (8), and 28 for hM4D- and mCherry-treated mice. (**G-H**) Relationship between fronto-parietal coordination during the consolidation phase and contextual fear memory retrievals. On days 1, 7, and 28, the freezing response to the CC was related either with concurrent PFC delta waves and RSC RIPc or concurrent PFC spindles and RSC RIPc (**H**). For mCherry-treated mice (black), the slope of a linear regression model reached significance (black lines) only on days 7 and 28 for concurrent PFC delta waves and RSC RIPc and concurrent PFC spindles and RSC RIPc, respectively. No relationship between the percentage of freezing and concurrent oscillatory patterns was observed in hM4D-treated mice (red).

In the results presented above, claustral inhibition disrupted both the consolidation of contextual memories and the co-occurrence of oscillatory events associated with episodic memory processes (*27*). Given the importance of PFC and RSC during the consolidation of remote memories (*35*), we investigated the relationship between claustrum dependent fronto-parietal coordination and the ability to retrieve a contextual fear memory. After contextual fear conditioning, fronto-parietal coordination was measured and the duration of freezing responses was quantified during conditioned context exposures on days 1, 7, and 28 (Fig. 2G). For control mice (Fig. 2H), we found linear relationships between freezing responses and concurrent retrosplenial RIPc and prefrontal delta waves (day 7) or spindles (day 28). No such relationships were observed in mice in which the claustrum was inhibited during the consolidation phase (Fig. 2H). These results demonstrate the relevance of fronto-parietal coordination for memory consolidation processes and their dependence on both the remoteness of the memory and on claustral activity.

To further investigate the neuronal mechanisms operated by the claustrum to coordinate fronto-parietal activity, we characterized neuronal spiking in PFC and RSC during the different oscillatory patterns. We observed that the activity of single units was modulated by the different oscillatory events (Fig. 3A-F), which was also noticeable at the population level (Fig. 3G-L). As previously described (*32*), neuronal spiking was first inhibited around the peaks of delta waves and then followed by a rebound of activity during the negative portion of the wave (Fig. 3G and J). The rebound of activity in prefrontal neurons overlapped with the onsets of RIPc in the RSC (∼ 200 ms after the delta peak; Fig. 3M). When *Vglut2*-positive claustral neurons were inhibited (confirmed by the drop in spiking activity recorded in the claustro-insular region; Fig. S6), the amplitude of both the peak and the rebound of delta waves were accentuated in PFC (Fig. 3J). These changes were accompanied by a more severe inhibition and a stronger rebound of spiking activity (Fig. 3G). Thus, claustral activity attenuated prefrontal delta waves and tamed neuronal activity fluctuations during these waves. Regarding spindles-related processes in the PFC (Fig. 3B, E, H, and K), neuronal spiking increased around the trough with the highest amplitude. However, when claustral neurons were inhibited, the peak activity was delayed roughly by 200 ms (Fig. 3H). Such delayed responses also occurred during concurrent RIPc in the RSC and spindles in the PFC with a lag of roughly 200 ms (Fig. 3N). Claustral inhibition affected neither the amplitude of RIPc in the RSC (Fig. 3L), nor the spiking response at the trough of the event (Fig. 3I) and the amplitude of spindles (Fig. 3K). In summary, claustral inhibition a) reduced the co-occurrence of prefrontal spindles and RIPc in the RSC (Fig. 1F, rightmost panel), and when these co-occurrences were observed, b) induced a time lag in their coordination.

**Figure 3.**
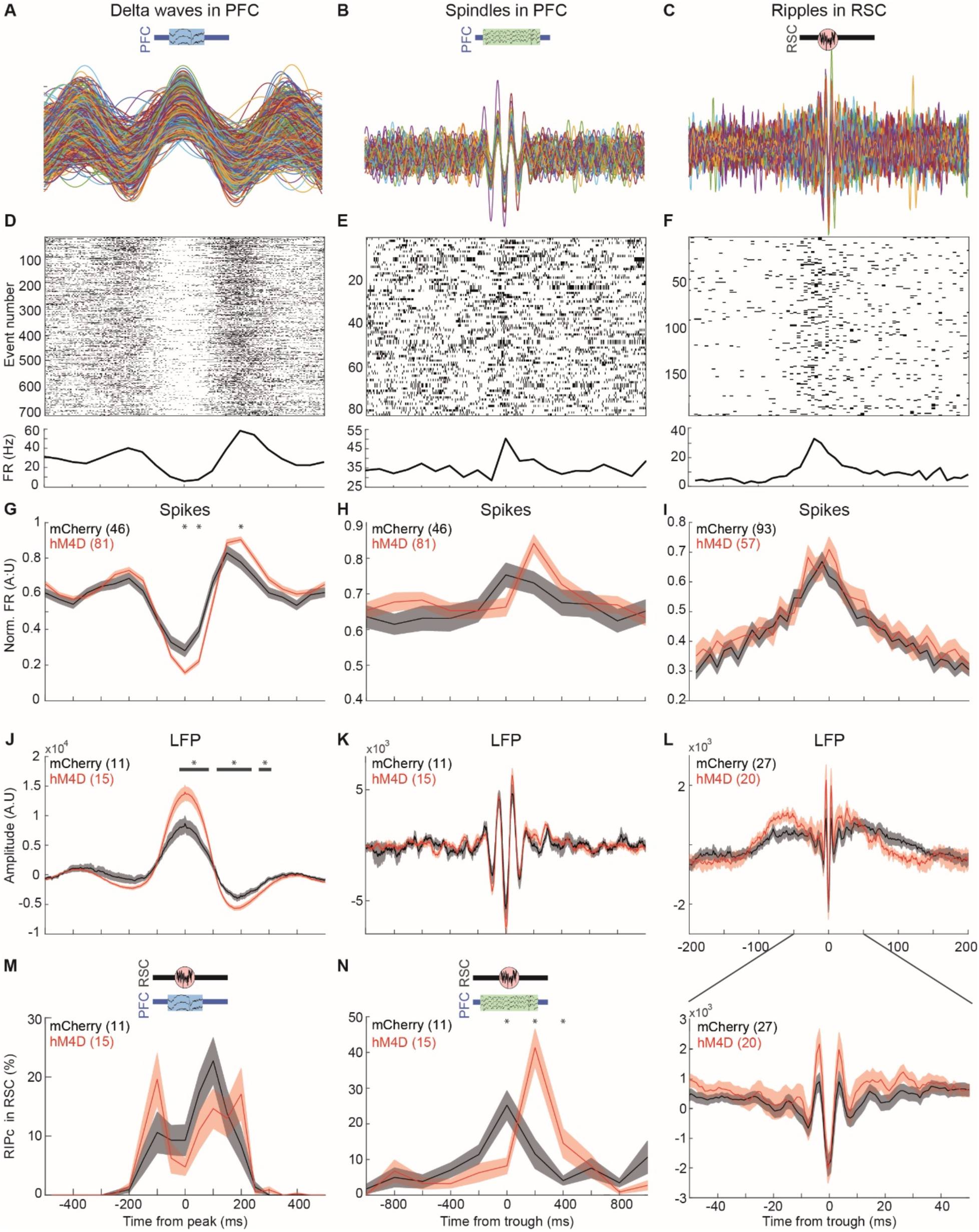
Neuronal spiking during concurrent retrosplenial ripples and prefrontal oscillatory patterns. (**A-C**) Examples of filtered LFPs for delta waves (**A**) and spindles (**B**) in the PFC and RIPc in the RSC (**C**) for a single recording site. (**D-F**) Representative examples of peri-event-time histograms and respective raster plots for a single unit recorded in the same tetrode than in (**A-C**). (**G-I**) Population activity of units during the oscillatory events in panels **J** to **L**. (**J-L**) Population waveforms of delta waves in the PFC (**J**), spindles in the PFC (**K**) and RIPc in the RSC (**L** and lower inset for a magnified view). (**M-N**) Distribution of RSC RIPc onsets referenced to the peak of the delta waves (**M**) and the trough of spindles in PFC (**N**). Data (shaded area; mean plus and minus the standard error of the mean; central line: mean) were collected in hM4D (red) and mCherry (black) treated mice. Significant differences are represented with either black stars or horizontal lines for each time bin when both p-values and FDR are < 0.05. If not specified, the scale of the x-axis is similar for all graphs in the same column.

Thus far, we showed that claustrum silencing leads to alterations of fronto-parietal communication relevant to memory consolidation processes. What about the ability to retrieve such memories? As the retrieval of remote fear memories is highly dependent on the PFC (*36, 37*) and on its projections to the hippocampus (*38*), we hypothesized that emphasizing signals from the PFC may promote such retrievals. We used two manipulations to bias fronto-parietal directional interactions. These manipulations consisted in inhibiting the excitability of either claustral projections to the RSC or claustral projections to the PFC. Using a retrograde viral cre delivery combined with local injections of cre-dependent hM4D DREADD in wild-type animals (as detailed above), claustrum projection neurons were inhibited prior to the retrieval of either a recent or a remote fear memory (Fig. 4A and B). When claustro-retrosplenial projections were inhibited, the freezing response to the conditioned context increased compared to the response of control animals (e.g., Movie S1 and S2). This effect was even accentuated for a remote memory (Fig. 4C top panel). This increase in the freezing response was not due to a general increase in immobility or other associative abilities because the freezing response to the conditioned cue was unaffected (Fig. 4C bottom panel). Such an enhancement of contextual memory retrievals did not occur when claustro-frontal projections were inhibited (Fig. 4C). To evaluate if contextual memory retrievals depended on signals from the PFC, we estimated the directional interactions between PFC and RSC during such retrievals (Fig. 4D). On day 28 after conditioning, when the memory enhancement was most accentuated, HFO signals from the PFC to the RSC were stronger in mice with inhibited claustro-retrosplenial projections compared to HFO signals in control mice (Fig. 4E). In summary, inhibiting claustral projections to the RSC emphasized signals from the PFC and enhanced the retrieval of remote contextual memories.

**Figure 4.**
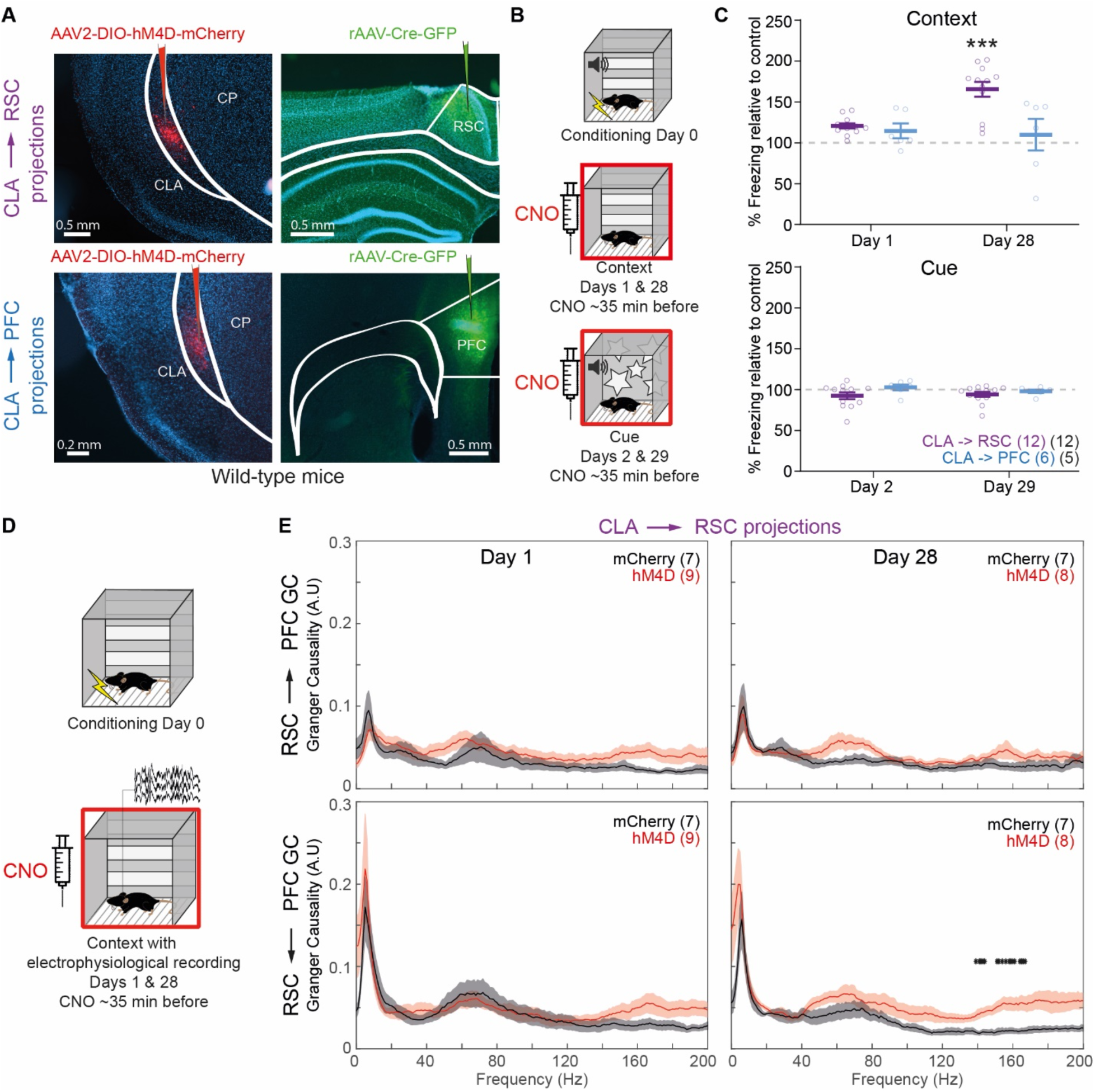
Clautral-dependent signals from prefrontal to parietal cortices facilitate remote contextual memory retrievals. **(A)** The claustro-insular region of wild-type mice (left panels) was infected with a cre-dependent viral construction containing either the inhibitory DREADD (hM4D) or not (mCherry). In the PFC or the RSC, a retrograde-transporting virus containing a cre-recombinase and associated with a Green Fluorescent Protein (GFP) was injected. (**B**) On days 1 (2) and 28 (29) after fear conditioning, mice were injected with CNO prior to measuring their freezing responses to the CC (CS). (**C**) Freezing response during re-exposures to the CC and the CS compared to control animals. (**D**) Wild-type mice with claustro-retrosplenial projections targeted using the cre-retrograde method (as above) were implanted with tetrodes in the PFC and the RSC. On days 1 and 28 post-conditioning, CNO was administered prior to contextual retrievals. (**E**) Directional interactions from RSC to PFC and from PFC to RSC during contextual retrievals. The number of pairs of tetrodes is shown in parenthesis and originated from 5 and 4 mice for hM4D and mCherry treatments, respectively. Significance was marked with a black star if both the p-value and the FDR were below 0.01.

## Discussion

The present study presents causal evidence on claustral-dependent fronto-parietal communication relevant to contextual memory processes. Some of these processes involved various oscillatory patterns, such as cortical ripples, delta waves, and spindles. These patterns participate in memory processes and are traditionally studied during sleep. However, these patterns and the consolidation of memories can occur outside of sleep periods (*39, 40*). In line with the emerging view that local oscillatory patterns are key signatures of memory processes, our results highlight the importance of their inter-areal coordination (*41*) and the role of the claustrum in this dynamic (*42*).

The claustrum, which connects almost the whole cortex (*8–10*) and is active during sleep (*17, 18, 43*), contains long-range glutamatergic projections preferentially found in supragranular layers of frontal, cingulate, and retrosplenial cortical areas; including layer 1 (*44*), which has been suggested as the storage location for long-term memories (*2*). Indeed, these properties place claustral activity as a major driving force for shaping interregional cortical dynamics relevant to memory processes. But the claustrum is not the only subcortical nucleus able to influence cortical interactions. For example, in vision, the pulvinar, a thalamic nucleus, facilitates directional interactions between visual areas (*45*). Another example is the reuniens, which modulates cortical coordination with the hippocampus during sleep (*46*). The medial septum also appears well suited to have such a property because of its role in both hippocampal theta rhythm and memory consolidation (*47*) and because it connects several cortical areas, including the retrosplenial cortex (*48*). Thus, it appears that several nuclei can compete or collaborate to modulate energy flow in the cortex (that is, i.e., traveling waves throughout the cortex). Maybe the claustrum’s unique projection pattern and dense connectivity (*9*) promote a preferential role in large-scale interactions with a bias toward associative areas. Thus far, no direct projections between the claustrum and the hippocampus have been described, but episodic memory processes are highly distributed and include multiple cortical areas in addition to the hippocampus (*3*). In the current study, it is most likely that the influence of the claustrum on fronto-parietal interactions reflects large-scale energy flows (*49*).

In conclusion, experimental and claustrum-dependent modulation of fronto-parietal communication had both an alteration and an enhancement of contextual memory processes. The underlying mechanisms probably involved shifts in neuronal response latencies rather than dramatic changes in the basal level of cortical activity. Given the widespread connectivity of the claustrum and the heterogeneity of its laminar projection profile (e.g., prefrontal versus entorhinal cortices), the precise influence of the claustrum on cortical interactions may differ according to the network involved. Nonetheless, the claustrum may be a critical cortical gatekeeper that targets and alleviates specific contextual-related psychophysiological symptoms (*50*).

## Acknowledgments

We thank Vladimir Kouskoff for his constructive comments on an earlier version of the manuscript. We also acknowledged all members of A.C. and I.R. laboratories for their participation in daily tasks to make scientific research possible.

## Funding

University of Geneva

The Swiss National Foundation (grant numbers: 31003A_172878 to A.C. and 310030E_135910 to I.R.)

The National Center of Competence in Research (NCCR) “SYNAPSY - The Synaptic Bases of Mental Diseases” The Swiss National Science Foundation (grant 51NF40-185897, A.C.)

The European Research Council (contract ERC-SyG-856439-CLAUSTROFUNCT to A.C. and I.R.)

The Divesa Foundation (A.C. and I.R.)

La fondation privée des HUG (A.C. and I.R.)

The Novartis foundation for medical research (A.C. and I.R.).

## Author contributions

Conceptualization: RFS, SM, JRR, AC, IR

Methodology: RFS, SM, JRR

Software: RFS, SM

Formal analysis: RFS, SM, JRR

Investigation: RFS, SM, JRR

Resources: AC, IR

Visualization: RFS, SM, JRR, AC

Funding acquisition: AC, IR

Project administration: AC, IR

Supervision: RFS, SM, AC, IR

Writing – original draft: RFS, SM

Writing – review & editing: RFS, SM, JRR, AC, IR

The authors declare no competing financial interest.

## Supplementary Materials

Materials and Methods

Supplementary Text

Figs. S1 to S6

References (51 to 60)

Movies S1 & S2.

## Data and materials availability

All data, code, and materials are available upon request.

## Materials and Methods

### Animals

Male and female *Slc17a6^tm2(cre)Lowl^*/J heterozygote mice (referred as *Vglut2*-cre in the text; the Jackson laboratory, strain number 016963). For some experiments, C57BL/6J mice originating from Charles River France were used. Mice (2-8 months old during the experiments) were housed in groups of 2–5, with the exception of animals implanted chronically with electrodes, which were single housed. Mice underwent a standard 12h light/dark cycle (7 am - 7 pm) in a 24°C temperature room with ad libitum access to food and water. Random treatment assignment was performed at the level of each cage using RFIDs. The matching of the treatment with RFIDs was unknown to the experimenters. All protocols involving alive mice are in accordance with the Swiss Federal Act on Animal Protection, the Swiss Animal Protection Ordinance and were approved by the University of Geneva and Geneva state ethics committees (authorization numbers: GE/94/20).

### Surgical procedures

Mice were anesthetized with isoflurane (Piramal Healthcare; 3–5% induction, 1–2% maintenance) before fixating their head to a stereotaxic apparatus (Stoelting). During the surgical procedure, artificial tears (Lacryvisc) were regularly applied onto the eyes and body temperature was maintained at 36° C using a rectal probe and a heating-pad (FHC) placed under the body. After two lateral and one medial subcutaneous injection of 25 µl carbostesin (AstraZeneca, 0.25 %) and dexamethasone (i.m. 20 µl at 4 mg/l), the skin overlaying the skull was either cut or, when a chronic implant is planned, removed. The anterior-posterior (AP) and medial-lateral (ML) axes of the surface of the skull were adjusted to form a horizontal plane. For viral injection, bilateral craniotomies were drilled dorso-ventrally (DV) to the claustrum (CLA) and a glass pipette containing the viral solution was inserted at two different coordinates: (1) +1.1 mm AP from Bregma, +/-2.85 mm ML from Bregma, -2.4 mm DV from brain surface and 100nL of viral solution; (2): +0.3 mm AP from Bregma, +/-3.2 mm ML from Bregma, -2.9 DV from brain surface and 75nL of viral solution. The coordinates for the prefrontal cortex were: +1.6 mm AP from Bregma, +/-0.3 mm ML from Bregma, -0.75 (1) and -1.75 (2) mm DV from brain surface and 100nL of viral solution at each site. The coordinates for the retrosplenial cortex were: -1.7 (1) and -2.7 (2) mm AP from Bregma, +/-0.4 mm ML from Bregma, -0.5 mm DV from brain surface and 100nL of viral solution at each site. In some experiment, the entorhinal cortex was targeted using the following coordinates: -0.2 mm AP from Bregma, +/-3.0 mm ML from Bregma, -3.0 mm DV from brain surface and 200nL of viral solution. The pipette was left in place for about 3 minutes before being removed. The following viruses (Addgene Co) were used: AAV2-*hSyn*-DIO-hm4D(Gi)-mCherry (hm4D), AAV2-*hSyn*-DIO-mCherry (mCherry), AAV2-*Ef1a*-DIO-Arch-EYFP (Arch), AAV2-*Ef1a*-DIO-EYFP (YFP), and pENN.AAV.*hSyn*.HI.eGFP-Cre.WPRE.SV40 (AAV retrograde). Then, the skin was sutured back to cover the whole skull except for mice planned for experiments involving optogenetic inhibition. In the latter case, mice were additionally implanted with 200 μm diameter optic fibers (OF; Thorlabs) glued (Precision Fiber Products, Inc) to ceramic ferrules (Precision Fiber Products, Inc). This procedure is fully described in (*51*). The OF-ferrule compounds were placed and cemented with a mixture of dental acrylic (Lang Dental) and glue at +1.1 mm AP from Bregma, +/- 2.9 mm ML from Bregma and at -1.9 mm DV from brain surface. To facilitate the recovery from the surgery, mice were given NaCl (s.c., 0.9% 500 µl), dexamethasone (i.m., 20 µl at 4 mg/l) and carprofen (s.c., 150 µl at 5 mg/l)

For experiment involving electrophysiological measurements, the animals recovered for 3-4 weeks after viral infection and underwent a second surgical procedure. As described above, mice were anesthetized, head-fixed and adjusted to expose their skull and to enable stereotaxic placement of electrodes. The electrophysiological implant consisted of 16 HFV-coated tungsten wires (0.0005”, California Fine Wire Company) connected to a 16-channel Omnetics electrode interface board (EIB, Neuralynx) and assembled as 4 tetrodes (*52, 53*). In addition, each tetrode was inserted into a polyimide tube (HVT Technologies) of either 2-3 mm (for cortical implantation) or 6 mm (for CLA implantation) to enable independent manipulation with a stereotaxic-fixed micro-forceps. In the right hemisphere, three small craniotomies were made to implant tetrodes in the prelimbic cortex (PFC, +1.6 mm AP from Bregma, +0.3 mm ML from Bregma, -0.75 mm DV from brain surface), in the retrosplenial cortex (RSP, -2.0 mm AP from Bregma, +0.5 mm ML from Bregma, -0.5 mm DV from brain surface) and in the CLA (+1.1 mm AP from Bregma, +2.85 mm ML from Bregma, -2.4 mm DV from brain surface). Animals had either a) 2 tetrodes in the RSP and two in the PFC or b) one in the CLA, two in the RSP and one in the PFC. Two stainless steel screws (PlasticOne) were driven into two additional craniotomies and used as electrophysiological ground and reference. Finally, the whole implant was cemented with dental acrylic.

### Behavioral experiments

During daytime, mice were daily habituated to handling and restraining procedures (for i.p. injections, optogenetic fibers plugging or headstage plugging for electrophysiological experiments) for three days prior to the start of the experiments. A camera placed above the home cages, the conditioning chambers, open field arenas and olfactory boxes recorded the animals’ behavior and was connected to the ANY-Maze software (Stoelting).

### Home cage activity and electrophysiological recordings

To evaluate the effects of claustral chemogenetic inhibition on cortical activity, electrophysiological signals were recorded during home cage behaviors. After being connected to the headstage, a mouse was placed back in its home cage. Note that the animal could freely move, rest or sleep and that the experimenter was outside of the room during the recordings. A first recording session started 10 min after a habituation period and lasted for 30 min. Then, either saline (0.4%) or clozapine N-oxide (CNO, TOCRIS Bioscience, 2 mg/kg) was injected i.p. and the second recording session started after 30 - 40 min and lasted for 30 min. Every other day the same procedure was repeated and the injected solution was alternated. Each mouse received a total of three saline and three CNO injections (but only data originating from CNO injections were reported in the present study). Only periods of immobility were further analyzed. These periods were automatically detected using ANY-Maze. The thresholds were adjusted manually on five randomly chosen videos to provide a stringent estimate of immobility. The thresholds (“on” and “off” thresholds were respectively 55 and 55, and the minimum freezing duration was set to 1 s) were applied to all videos from all mice. This method was preferred to the ANY-Maze built-in immobility evaluation because the headstage wires movement prevented the latter procedure to work. In a few cases, the field of view had to be re-adjusted so that the headstage wires were less visible. Immobility was binned in segments of 2 s and expressed as the total amount of time in each bin.

### Fear conditioning

Custom-made square (20 x 20 x 20 cm) or octagonal (10cm each side, 20 cm high) Plexiglas chambers displaying a black & white pattern on its walls and containing a metallic grid floor were used. The chambers were placed in a soundproof box containing two speakers and two LEDs. The ANY-Maze software was also used as an interface to control sound, electrical shock and laser beam emission. During fear conditioning, the animal was placed in the center of the chamber and a 180 s period of acclimatization started. In experiments where a conditioned stimulus (CS) was present, the CS (30 s white noise at 80 dB or 3000Hz tone at 80 dB in the case of 2^nd^ conditionings) was then presented and co-terminated with an unconditioned stimulus (US). This US consisted of a 2 s electrical shock (0.5mA) delivered through the metallic grid. The CS-US pairing was presented three times in total (30 s inter-sounds). During contextual fear conditioning (no CS), the US was delivered right after the acclimatization period, then two additional times (1 min inter US). For optogenetic inhibition during fear conditioning, Arch infected mice received one pulse of laser beam (550 nm orange light, 11-14mW measured at the tip of the OF before implantation) either paired with the electrical shock (starting 2 s before and co-terminating with the shock) or co-starting with the CS and lasting 4 s. The conditioned context (CC) was characterized by the location of the soundproof box in the room, the shape of the chamber (square or octagonal), the specific black & white pattern on the walls (e.g., stripes, gears, stars and more), the LEDs (either white or green), the cleaning solution of the chamber (either 70% ethanol or a commercial 38% bio-ethanol (Oecoplan) and the grid on the floor (either metallic or plastic).

Following conditioning (day 0), fear memory recalls (300 s) for the CC occurred on the next day (day 1), on the next week (day 7) and, in some experiments, mice underwent an extinction procedure (see next paragraph for details) after four weeks (day 28) and the first 150 s of this procedure were used as a context recall. These recalls consisted in placing the mouse in the same CC than during conditioning but without the CS. Memory recalls for the CS occurred two days (day 2), one week and one day (day 8) after conditioning. These recalls consisted in placing the mouse in a different sound-proof box, a chamber displaying different black & white patterns on its walls, LEDs with different colors, different cleaning solution and a different floor grid (in plastic) than during conditioning. After 180 s, the CS was presented in a similar protocol as during the conditioning.

One week after the last recall, some mice, which were planned to undergo a second conditioning (as control experiments), underwent an extinction procedure. This procedure consisted in placing the mouse in the CC and, after 150 s, in repeating the presentation of the CS 20 times (30 s inter-CS). On the next day, an extinction test consisted in placing the mouse in the CC and repeating the CS three times in a similar timeline than during the conditioning but without the US. This procedure ensured that both groups of mice reached a comparable level of memory extinction.

To quantify the ability of the animal to retrieve a fear memory, a traditionally used conditioned response (CR) of the animals, the duration of a freezing state, was estimated in a semi-automatic manner. First, a manual scoring for each animal was performed off-line by an observer (naïve for the treatment) for one CC and one CS recall. Then, the parameters in ANY-Maze software were tuned to match the score measured by the observer. In details, the freezing thresholds (referred to as ‘on’ and ‘off’ in ANY-Maze) were set as equal and were tuned to match the total amount of freezing time obtained by manual scoring (the minimum freezing duration in ANY-Maze software was set at 2 s). This procedure was applied separately for each mouse and for each type of recall (CC and CS) and then the thresholds were applied to the other days with similar type of recalls. The advantage of this semi-automatic procedure to the automatic one implemented in ANY-Maze is a more precise estimation of the duration of the CR. For retrievals during which electrophysiological activity was recorded, freezing was obtained by manual scoring only, due to the presence of a wire that interfered with automatic freezing estimation.

Overall, five animals developed an escaping strategy during fear retrievals. This was expressed as a jumping response. The animals were excluded from the analyses.

### Optogenetic and chemogenetic manipulations

Optogenetic inhibition was achieved using a cre-dependent expression of the Archaerhodopsin (*54*) in *Vglut2*^+^ claustral neurons and by delivering an orange light (550 nm, 14 mW at the tip of the fiber) through the implanted optic fibers using a laser (Doric Lenses). Chemogenetic inhibition, based on Designer Receptors Exclusively Activated by Designer Drugs (DREADD; (*55*)), was used to elucidate the role of the CLA during memory consolidation, during locomotion and quietness. Mice expressing either hM4D or mCherry were injected (i.p.) with CNO (2 mg/kg). For mice that underwent fear conditioning, the first injection occurred after removing the animal out of the conditioning box. Once in the home cage, the injections were repeated every two hours for a total of 4 injections per mouse during daytime. Mice were placed individually in clean cages and their movement were tracked for 3h30 to evaluate the effects of CNO on home cage behaviors. To further characterize CLA chemogenetic manipulations on locomotor activity and once an animal terminated all of the fear conditioning experiments described above, CNO was injected (i.p.) 30 – 40 min prior to monitoring the distance travelled in an open field arena (46 x 46 cm, white walls and floor for 10 min. The mouse position was tracked using the ANY-Maze software after placing the animal in the center.

### Memory consolidation period

A couple of days prior to the conditioning, mice with a chronic electrophysiological implant were habituated to a clear Plexiglas box (20 x 20 x 40 cm) containing clean bedding. This box was connected to an odor delivery system that could diffuse odors. After fear conditioning, mice were placed in the box, and after a habituation period (3-5 min depending on the stillness of the mouse), either the cleaning solution used during fear conditioning or an ethyl butyrate solution were peuso-randomly diffused for 10 s (30s ITI, 10 repetitions per odor). Then, CNO was injected i.p., the mouse was placed back in the box and could rest for 30 - 40 min before the same odor diffusion protocol was repeated. Electrophysiological signals were recorded during the whole time except during the i.p. injection and the following resting period. As an initial analysis did not reveal any evoked response neither in the LFP nor in the firing rate, the effect of diffused odors on neuronal activity was not be further pursued.

### Electrophysiological recordings

A HS-16 headstage (Neuralynx) connected the animal’s EIB to a Digital lynx recording system. The mouse and its recording cages were put on a pressurized table (Co) to reduce movement artefacts. No apparent discomfort or movement anomaly was noticeable.

### Signal processing

Broadband signals were band-pass filtered (0.1 Hz to 9 kHz), digitalized at 32 KHz sampling rate and synchronized with videos of the animals via the Cheetah Software (Neuralynx). A detailed description of the procedure to extract Local field potentials (LFPs) and unit activity is found in (*56*) and in (*57*), respectively. Shortly, local field potentials (0.1-600 Hz for spectral quantities or 0.1-1250 Hz for the detection of oscillatory events) underwent a linear detrend and a z-score normalization using custom-made Matlab (Mathworks) scripts. Using a freely available package referred to as NDManager (http://neurosuite.sourceforge.net), unit activity and spike waveforms were extracted by applying a threshold to the high-pass filtered signal (spike detection threshold = 1.3; peak search window = 20; samples per waveform = 40; peak sample = 20). Spike waveforms were clustered using the freely available python-based package Klustakwik and single units were isolated from multi-unit activity either with Klusters (included in NDManager) when data sets exceeded 5 min or with MClust (Matlab based) for smaller data sets. Klusters is more reactive than MClust for big data sets but we implemented a routine to handle spike overlaps in MClust (*58*), which is absent in Klusters. A cluster was considered as a single unit (SUA) if 1) a minimum refractory period of 1-2 ms was present in the autocorrelogram, 2) the cluster was clearly separated from all of the surrounded clusters in at least one dimension, 3) no apparent cross-correlation with the other clusters and 4) the spikes inside the cluster had a stereotyped waveform. All clusters that did not satisfy these criteria and that did not consist of artifacts were grouped as one cluster of multi-units.

### Spectral analyses

Spectral analyses of LFPs were performed using the Matlab-based toolbox named Chronux (*59*). Time-frequency power spectra were obtained using nine tapers, a time-bandwidth of five and using non-overlapping windows of 2s. For each tetrode, the median power (or any other quantity) between the four corresponding electrodes was used. To identify time-windows contaminated with artefacts, we used an automated Kurtosis-based artefact detection (*56*). Shortly, noisy 2s-windows (with extreme high power) were excluded until the Kurtosis values of the density distribution of the other time-windows was equal or lower to 4.0 at all frequencies. This method yielded the removal of 5 to 15 % of all time-windows. Only tetrodes with identified units were used in this study.

Directional Wiener-Granger causalities (GC) were quantified using the Matlab-based toolbox named MVGC Multivariate Granger Causality (*60*). The Akaike information criterion was used to choose the model order (maximum model order of 100) for each 2s-time-window. The 25^th^, 50^th^ and 75^th^ percentiles of the model orders were 20 (33 ms), 26 (43 ms) and 34 (57 ms), respectively. All other parameters were set to the defaults. 2s-time-windows were automatically checked for stationarity, collinearity and other basic requirements using built-in functions and were excluded otherwise. Frequency resolution was set to 1Hz.

### Data and frequency band selection

The following analyses only include period of immobility. The reason is that to analyze periods of activity would require to characterize the different behaviors and to compare each one of them separately. The first and simplest approach was to start with one simple behavior: immobility. These periods consisted in 2s-bins having 2 s of immobility. Data from a recording session were included only if the total number of bins when the animal was considered as immobile was 30 (equivalent to 1 min) or more. The reason for this criterion was that, after visual inspection, the automated detection of oscillatory events (see below) yielded too many false positives with parameters that worked well for data sets with a large number of immobility bins. At the population level, the 25^th^, 50^th^ and 75^th^ percentiles of the number of bins with immobility per 30 min session were 114 (228 s), 339 (678 s) and 586 (1172 s), respectively. The median spectral value within a specific frequency band (delta 0-4 Hz, theta 4-10 Hz, alpha 10-20, beta 20-30 Hz, gamma 30-120 Hz and ripple-related 120-250 Hz) was representative of that band.

### Detection of oscillatory events

The detection of high-frequency ripple-like oscillations (RIPc), delta waves and spindles were performed using the Matlab-based Freely Moving Animal toolbox (https://fmatoolbox.sourceforge.net). The procedure consisted in filtering LFPs into a narrow-band signal corresponding to the operating frequency of the specific event. Then, several amplitude thresholds and constrains on the minimum and maximum durations of the events were applied. RIPc (100-250 Hz) were identified with thresholds at 4.0 and 6.0 folds of the standard deviations for the beginning/end and peak, with a minimum inter-RIPc interval of 30 ms, minimum duration of 30 ms and a maximum duration of 110 ms. Delta waves (0-6 Hz) were identified with thresholds at 1.0/2.0 and 0.0/1.5 folds of the standard deviation for minimum/maximum peaks and troughs, respectively. The minimum and maximum durations were 160 ms and 500 ms. Spindles (9-17 Hz) were detected with a minimum envelope tail of 3.0 (z-score value), a minimum envelope peak of 6.0 (z-score value), minimum and maximum durations of 750 ms and 3500 ms, respectively. All of the parameters presented above were modified from defaults values to reduce the number of false positives (estimated visually) on randomly chosen data sets. These detections were applied to each electrode of one tetrode. As the signals within one tetrode were very similar, the averages (start, end and amplitude of the event) between the different electrodes were used to represent the tetrode. The only exceptions were the start and end of RIPc where the earlier onset and latest offset between the electrodes were used. Finally, data originated from tetrodes with at least 50 events (RIPc, delta waves and spindles) for one session were included in the analyses.

### Histology

To mark the electrodes positions, electrolytic lesions were induced using a NanoZ device (White Matter LLC). A 12 μA current was applied to each electrode for 8 s, twice. To verify the specificity of viral infections, mice were euthanized with an injection of pentobarbital (200 mg/kg) and phosphate-buffered saline (20 ml; 0.1 M; PBS) was perfused transcardially (20 ml) followed by a 4% paraformaldehyde solution (50 ml; 0.1 M in PBS; PFA). If an implant was present, it was carefully removed before extracting the brains. The brains were stored in PFA overnight (12-24 hrs) and then kept in PBS. Brains were embedded in 4% agarose before being cut into 40 um coronal slices with a vibratome (Leica VT S10000) and stored in 0.1% PBS Sodium Azide. The slices were pre-incubated in PBS 1% BSA 0.3% triton for one hour at room temperature. First antibody (rabbit anti-GFP INVITROGEN A6455 1:2500; chicken anti-mCherry ABCAM ab205402 1:2000) was diluted in pre-incubation buffer and applied to the slices for overnight at 4°C. The slices were washed for 3 times 10 minutes with PBS. Secondary antibody (donkey anti-rabbit alexaFluor 488 INVITROGEN A-21206 1:1000; goat anti-chicken alexaFluor 568 INVITROGEN A-11041 1:1000) was diluted in PBS 1% BSA and applied to the slices for one hour in the dark at room temperature. The slices were washed for 2 times 10 minutes with PBS. For cell nuclei staining, Hoechst (INVITROGEN H3570 1:5000) was diluted in PBS and applied for 20 minutes in the dark at room temperature. Slices were mounted onto glass slides and cover-slipped with Immu-Mount, (THERMO 9990412). Every third slice was mounted for visual inspection under the microscope.

Prior to any data analysis, the exclusion of animals based on histological verification was performed. Animals were excluded if the electrode locations were misplaced or if the locations of the infections failed. An agreement between at least two out of three experimenters was reached before excluding an animal. Each experimenter inspected the brain slices independently before sharing his decision on animal exclusions.

### Statistical analyses

To test the effect of claustral Vglut2 inhibition on cortical interaction, mice received several injections CNO. For LFP analyses, the measured quantities originating from the repeated injections of the same drug were averaged to avoid statistical dependence. For multiple comparisons, the false discovery rate was used if multiple comparisons were performed. Statistical analyses were performed using custom-written MATLAB scripts. GraphPad PRISM was used to analyzed behavioral data.

### Supplementary Text

All datasets for behavioral results (Fig. 2E-F and 4C) consist in replicates from original pilot studies. The data of the pilot studies were not merged with the datasets used in the present study.

Additional text and statistical information for Figures:

Figure 1:

**(A)** (B) In the inset, the thickness of the spectra represents the standard error of the mean.

**(B)** For different frequency bands, each GC measure (between one tetrode in PFC and another tetrode in RSC) was represented by the median GC value within the frequency band of interest. To level the different frequency bands, each GC measure was normalized by the maximum population-averaged value (two directions and two treatments). A three way ANOVA (treatment x direction x frequency band) yielded significant effects of treatment (F(1) = 19.76, p = 0.00001) and frequency band (F(5) = 27.23, p = 3*10^-24^) and an interaction between treatment and direction (F(1) = 7.07, p = 0.0081). No effect of direction (F(1) = 1.59, p = 0.206) and no interaction between treatment and frequency band (F(5) = 1.86, p = 0.98) and between direction and frequency bands (F(5) = 0.87, p = 0.49) were found.

**(C)** To quantify the difference between the two GC directions, the GC measures for the retrosplenial–to-prefrontal cortex (RSC->PFC) were subtracted from the GC values for prefrontal-to-retrosplenial cortex (PFC->RSC), and these differential GC values were divided by the mean between the two directions. Comparisons between treatments were performed using t-tests with a false discovery rate correction. The respective p-values for the different comparisons were 0.172, 0.216, 0.679, 0.588, 0.002 and 0.012 (from left to right and from top to bottom panels). We also determined whether each population was significantly different from zero using t-tests with a false discovery rate. The p-values for the different comparisons were 0.6 (0.00375), 0.215 (0.681), 0.951 (0.338), 0.726 (0.0232), 0.339 (0.000028) and 0.274 (0.0035) (from left to right and from top to bottom panels), for the mCherry (hM4D) groups, respectively.

Error bars represent the mean and the standard errors of the mean.

(**F**) For RIPc in the PFC and in the RSC, the sequential occurrence is shown separately in the 1^st^ and 4^th^ panels. The 2^nd^ and 3^rd^ panels correspond to the co-occurrence of RIPc in the PFC and either delta waves or spindles in RSC, respectively. The 5^th^ and 6^th^ panels show data for concurrent RIPc in the RSC and either delta waves or spindles in PFC, respectively. Comparisons between treatments were performed using t-tests. The p-values from left to right panels are 0.996, 0.224, 0.562, 0.337, 0.033 and 0.00003.

Figure 2:

**(A)** Optic fibers were implanted above the CLA. In the conditioned context (CC), the laser pulse (550 nm, 14 mW at the tip for 4 s.) was delivered during the last 4 s. of the conditioned stimulus (CS; White noise at 80 dB for 30 s.), which was paired with the electric shock during the last 2 s of the sound.

(**C-D**) CNO (2mg/kg, i.p) was injected right after conditioning, then every 2.0 hours for a total of four injections. During context retrievals, a two-way ANOVA with repeated measures yielded a significant effect of treatment (F(1) = 17.60, p = 0.0003), of days (F(2) = 9.992, p = 0.0002) and their interaction (F(2) = 5.392, p = 0.0079). Post-hoc comparisons with a Bonferroni correction showed a significant difference in the level of freezing between YFP and Arch treated mice on days 7 (p=0.0441*) and 28 (p<0.0001****), but not on day 1 (p = 0.1244). For the freezing response to the CS, a two-way ANOVA with repeated measures yielded neither an effect of treatment (F(1) =0.128, p=0.7238), of days (F(1)=0.000259, p=0.9873) nor their interactions (F(1)=0.3291, p=0.5717).

(**G-H**) First, mice underwent contextual fear conditioning, then received one injection of CNO, and fronto-parietal activity was recorded during the consolidation phase.

**(G-H**) During the context retrieval, a two-way ANOVA with repeated measures yielded a significant effect of treatment (F(1) = 23.78, p < 0.0001) and of days (F(2) = 4.836, p = 0.0115). Their interaction was not significant (F(2) = 1.017, p = 0.3681). Post-hoc comparisons with a Bonferroni correction showed a significant difference in the level of freezing between mCherry and hM4D treated mice on days 7 (p=0.0043**) and 28 (p=0.0005***), but not on day 1 (p = 0.1169). For the control experiment with saline injections instead of CNO, a two-way ANOVA with repeated measures yielded a significant effect of days (F(2) = 13.40, p < 0.0001), but not of treatment (F(1) = 0.001529, p = 0.9691) or their interaction (F(2) = 0.1771, p = 0.8381). For simplification, only values for day 28 are presented for the control experiment. For the freezing response to the CS, a two-way ANOVA with repeated measures yielded neither an effect of treatment (F(1)=0.4253, p=0.5196), of days (F(1)=1.607, p=0.2154) nor their interactions (F(1)=0.7687, p=0.3881).

(**J**) The relationship between concurrent RSC RIPc and either delta waves or spindles in PFC was assessed with a linear regression model. For mCherry (hM4D) populations (left panels), the p-values for a slope different from zero were, from top to bottom, 0.192 (0.986), 0.0037 (0.741) and 0.172 (0.694). For mCherry (hM4D) populations (right panels), the p-values were, from top to bottom, 0.76 (0.991), 0.57 (0.387) and 0.0002 (0.484).

Significance levels are represented with stars as follow: p<0.05 (*), p<0.01 (**), p<0.001 (***) and p<0.0001 (****).

Figure 3:

(**A-C**) In (**A**), the traces were aligned according to the highest peak of each event. In (**B**) and (**C**), the events were aligned according to the most negative troughs.

(**D-F**) For visualization purposes, the bin size was set to 5 ms and the threshold for the color scale in the raster plots was set at 1 spike.

(**G-I**) Each peri-event-time histogram was divided by its maximum firing rate to compensate for differences in baseline activity (see Fig. S6 for the same data set without normalization). Comparisons between treatments were performed using t-tests for each bin and significance was marked if both the p-values and the false discovery rates were below 0.05.

(**J-L**) For each recorded site, the local field potentials were averaged to represent the oscillatory events of that site.

Error bars represent the mean and the standard errors of the mean. Shaded areas represent the standard errors of the mean.

Figure 4:

(**C**) For each experimental animal, its percentage of time spent freezing was divided by the average freezing time of all control animals (cage mates) tested the same day. However, comparisons were performed on the percentage freezing time between experimental groups and their respective controls. Significance was marked with black stars if the p-value, Bonferroni-corrected, was below 0.0001 (***). For the freezing responses to the CC (left panels) in the CLA->RSC group, a two-way ANOVA with repeated measures yielded a significant effect of treatment (F(1) = 16.99, p = 0.0004), days (F(1) = 9.242, p = 0.006) and their interaction (F(1) = 5.307, p = 0.0311). Post-hoc comparisons with a Bonferroni correction showed a significant difference in the level of freezing between hM4D and mCherry treated mice on day 28 (p<0.0001****). In the CLA->PFC group, a two-way ANOVA with repeated measures yielded no significant effect of treatment (F(1) = 0.71, p = 0.42), days (F(1) = 0.1618, p = 0.6969) or their interaction (F(1) = 0.03839, p = 0.849). For the freezing response to the CS (right panels) in the CLA->RSC group, a two-way ANOVA with repeated measures yielded no effect of treatment (F(1) = 4.211, p=0.0523), or the interaction between treatment and days (F(1)=0.019, p=0.8912), but yielded a significant effect of days (F(1)=16.53, p=0.0005). In the CLA->PFC group: a two-way ANOVA with repeated measures yielded no effect of treatment (F(1) = 0.008, p=0.9306), or the interaction between treatment and days (F(1)=0.997, p=0.3441), but yielded a significant effect of days (F(1) = 9.245, p=0.014).

(**E**) Group differences were tested (t-test) for each frequency, each day and each direction.

Error bars represent the mean and the standard errors of the mean. Shaded areas represent the standard errors of the mean.

**Fig. S1.**
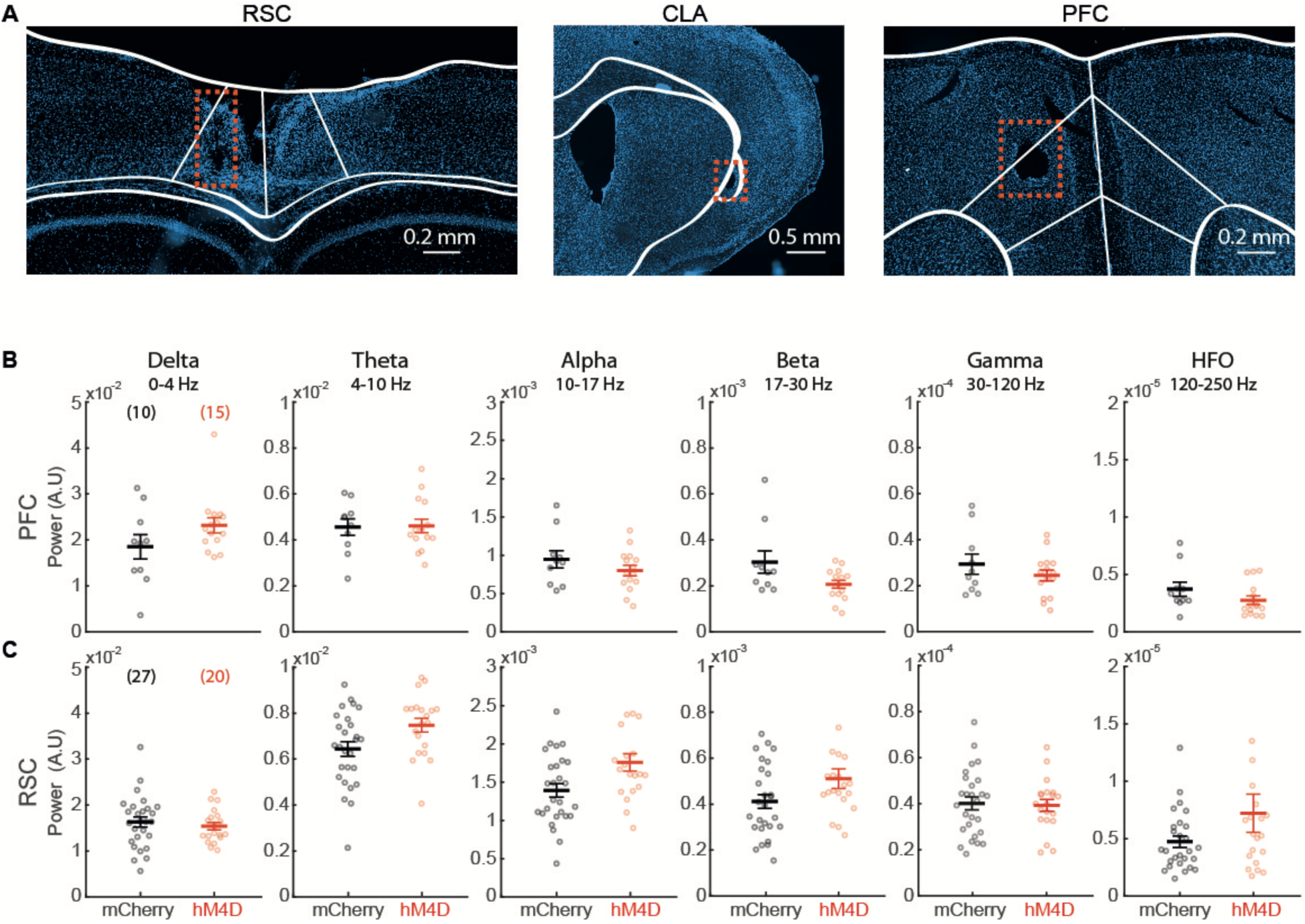
Claustrum inhibition does not alter the amplitude of neuronal oscillations in PFC and RSC. (**A**) Tetrodes were placed systematically placed both in the prefrontal cortex (PFC) and in the retrosplenial cortex (RSC) and, in some mice, also in the claustrum (CLA). Electrolytic lesions were performed to verify electrode locations (dashed red boxes). (**B**-**C**) Power of local field potentials for different frequency bands shown separately for the PFC (**B**) and RSC (**C**). Data are presented separately for LFPs recorded in hM4D (red) or mCherry (black) treated mice. Comparisons were performed using t-tests with a false discovery rate correction. The respective p-values for the different comparisons were 0.128 (0.542), 0.903 (0.154), 0250 (0.152), 0.155 (0.058), 0.297 (0.839) and 0.168 (0.116) for the Delta, Theta, Alpha, Beta, Gamma and HFO in the PFC (RSC), respectively. Error bars represent the mean and the standard errors of the mean.

**Fig. S2.**
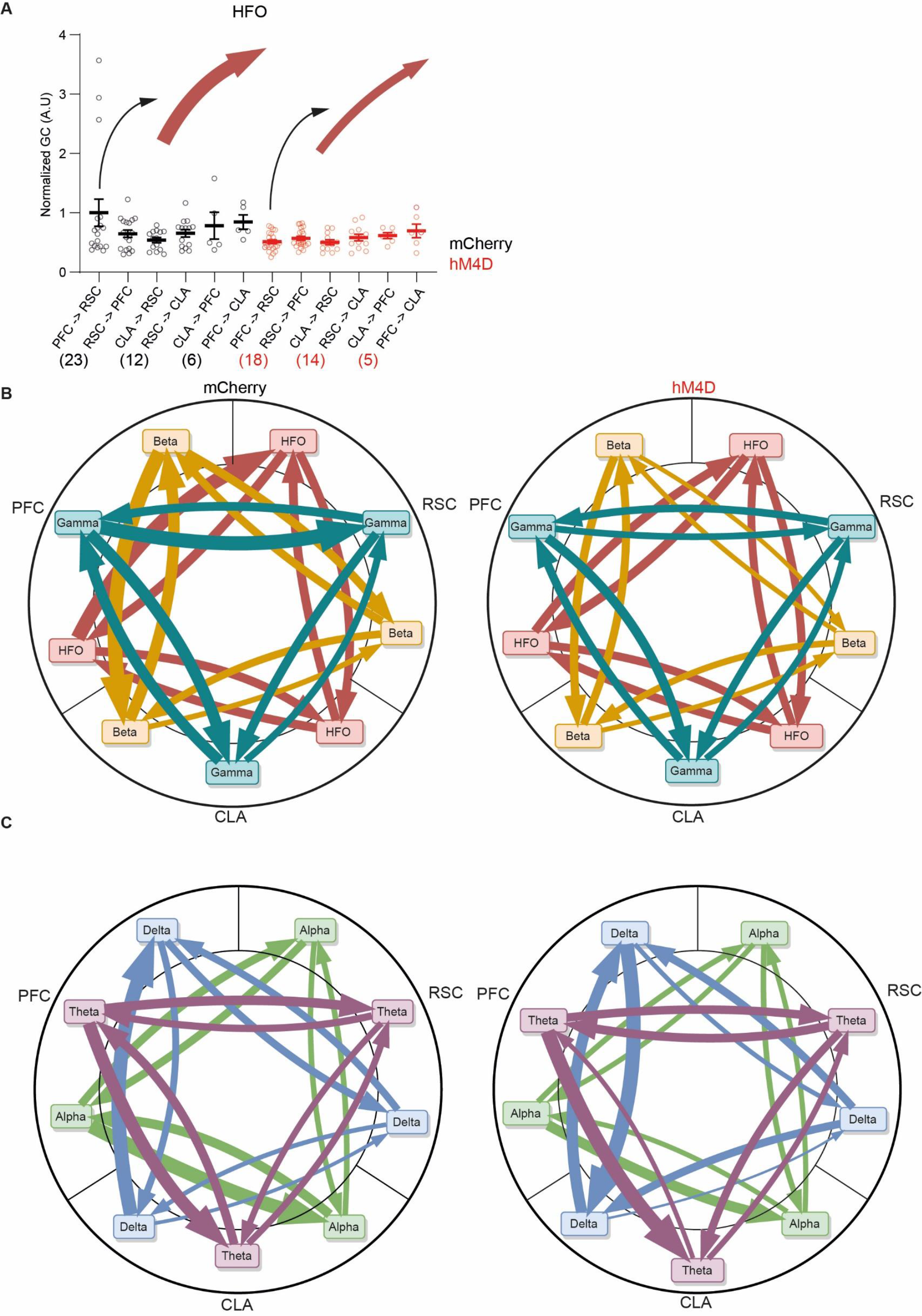
Directional interactions in the claustro-cortical and in the fronto-parietal networks. (**A**) Example of a scatter plot of Granger causality measures in the gamma frequency band. Mean and standard errors of the mean are presented. The width of the arrows represent the mean of the population as in **B** and **C**. (**B-C**) For different frequency bands (Delta 0-4 Hz, Theta 4-10 Hz, Alpha 10-17 Hz, Beta 17-30 Hz, Gamma 30-120 Hz, high-frequency oscillations (HFO) 120-250 Hz), each GC measure between inter-areal pairs of tetrodes (either in PFC, in RSC or in the claustro-insular region (CLA)) was represented by the median GC value within the frequency band of interest. To level the different frequency bands at a comparable amplitude, each GC measure was normalized by the maximum population-averaged value (two directions and two treatments). The width of the arrows represents the mean value of the population (n = 23 (18), 12 (14) and 6 (5) for PFC-RSC, CLA-RSC and PFC-CLA pairs of tetrodes collected in 11 (9) mice with hM4D (mCherry) treatments, respectively). Double arrows represent the maximum GC value across all treatments/directions and dashed lines represent its minimum. A three-way ANOVA (treatment x direction x frequency band) yielded significant effects of treatment (F(1) = 17.35, p = 0.00003), frequency band (F(5) = 23.31, p = 10^-21^) and direction (F(5) = 14.26, p = 10^-13^). It also revealed significant interactions between treatment and direction (F(5) = 3.94, p = 0.0015), between treatment and frequency band (F(5) = 3.44, p = 0.0043) and between direction and frequency band (F(25) = 1.96, p = 0.0034). High (**B**) and low (**C**) frequency bands were presented separately for visualization purposes.

**Fig. S3.**
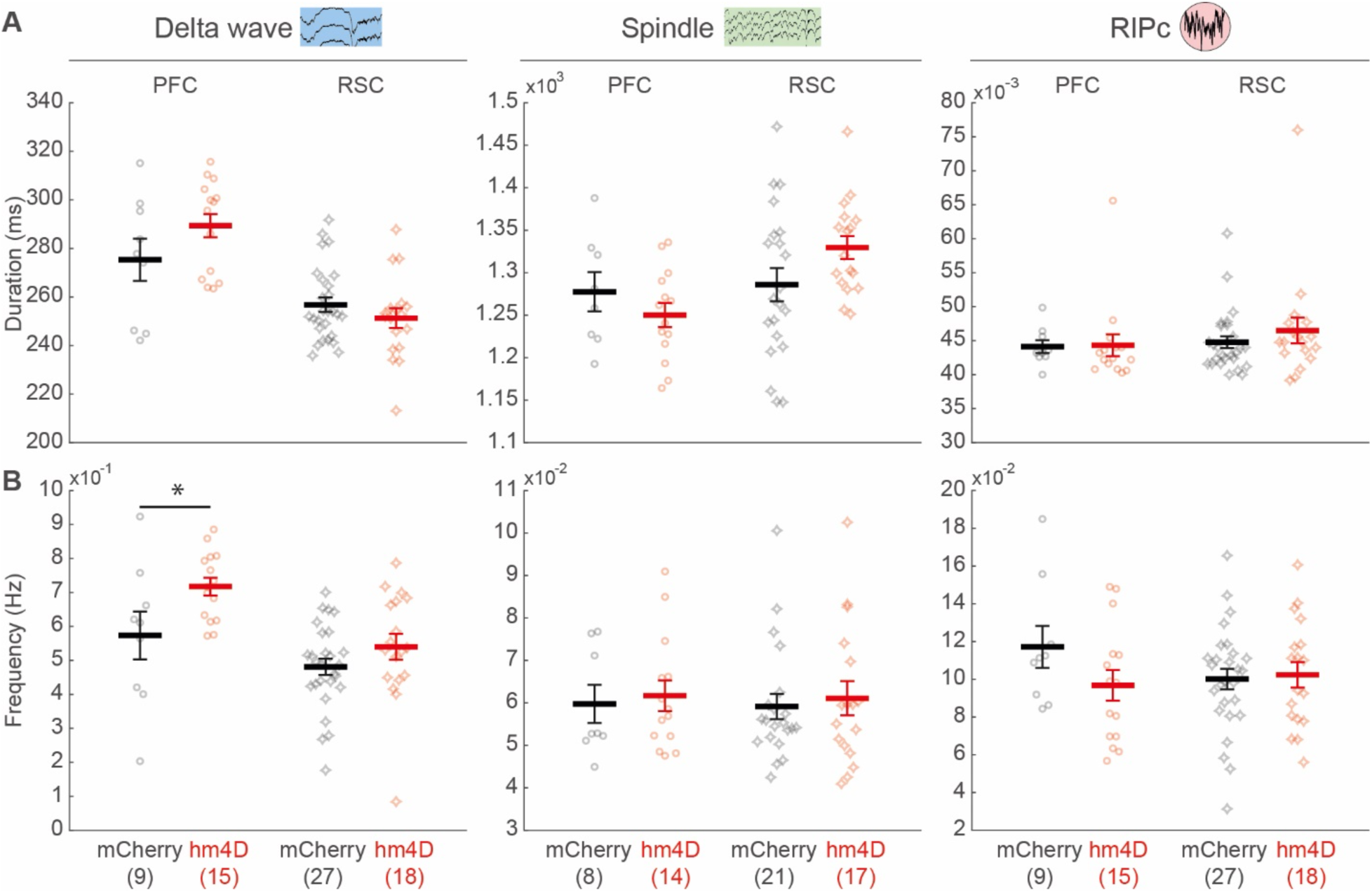
Impact of claustral inhibition on various oscillatory patterns in the fronto-parietal network. Durations (**A**) and frequency (**B**) of delta waves (left panels), spindles (middle panels) and high-frequency events (RIPc; right panels) in PFC and in RSC. Statistical comparisons between treatments were evaluated using t-tests with a False Discovery rate. The p-values for the PFC (RSC) groups were, from left to right and from top to bottom, 0.138 (0.268), 0.3007 (0.088), 0.934 (0.363), 0.034 (0.173), 0.749 (0.701) and 0.145 (0.796). Error bars represent the mean and the standard errors of the mean.

**Fig. S4.**
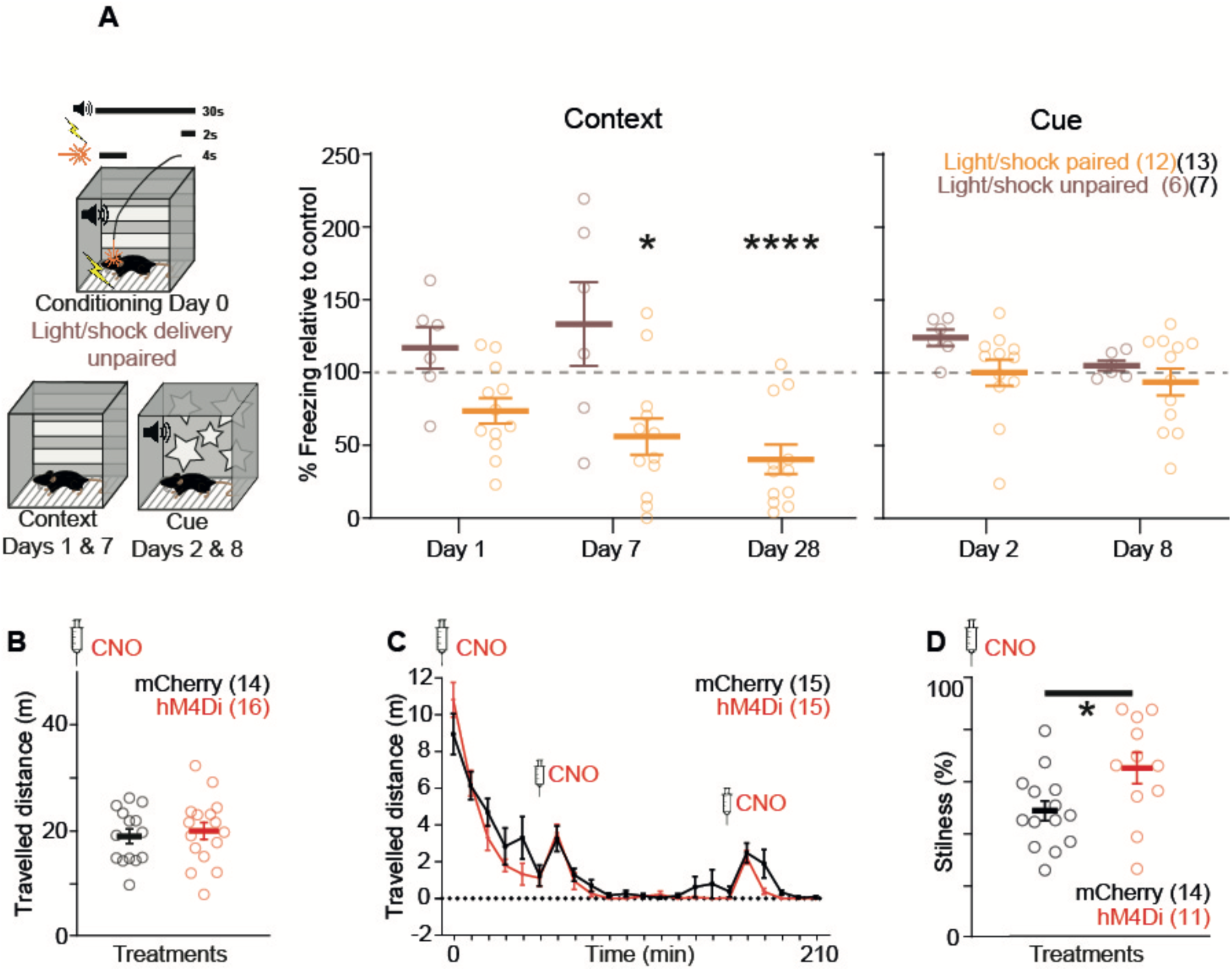
Supplementary behavioral experiments to characterize the consequences of claustral inhibition. (**A**) Unpaired claustral inhibition and electric shock delivery during fear conditioning. In the conditioned context (CC), the laser pulse (550 nm, 14 mW for 4 s.) was delivered at the beginning of the conditioned stimulus (CS; Pure tone at 3000 kHz, 80 dB for 30 s.), which was paired with the electric shock during the last 2 s of the sound. The freezing response to the CC (CS) were measured on days 1 (2) and 7 (8) post-conditioning for mice infected with a virus containing either an inhibitory opsin (Arch, n=6) or with a virus lacking the opsin (YFP, n=7). For each experimental animal, its percentage of time spent freezing was divided by the average freezing time of all control animals (cage mates) tested the same day. However, comparisons were performed on the percentage freezing time between experimental groups and their respective controls. For the CC, a two-way repeated measures ANOVA (treatment x days) yielded no significant effect of days (F(1) = 3.29, p = 0.097), treatment (F(1) =1.401, p = 0.2615), or interaction between treatment and days (F(1) = 0.07659, p = 0.7871). For the CS, a two-way repeated measures ANOVA (treatment x day) yielded a significant effect of days (F(1) = 18.27, p = 0.0013), but no significant effect of treatment (F(1) = 1.317, p = 0.2755) or interaction between treatment and days (F(1) = 3.818, p = 0.0766). (**B**) Total distance travelled in an open field arena 30 – 40 min after an injection of CNO in hM4D (n=16) or mCherry (n=14) treated mice. A Mann-Whitney test yielded no significance (Z = ‒0.39 p = 0.69). (**C**) Total distance travelled in a clean homecage following three injections of CNO at 0, 50 and 160 minutes. Animals were single housed for this test (hM4D, n=15; mCherry, n=15). A two-way repeated measures ANOVA (treatment x time bin) yielded a significant effect of time bins (F(21) = 28.31, p < 0.0001), but no significant effect of treatment (F(1) = 1.571, p = 0.2246) or interaction between treatment and time bins (F(21) = 1.41, p = 0.1079). (**D**) Effect of claustral inhibition on the percentage of time spent still. hM4D (n=11) or mCherry (n=11) treated mice were injected with CNO and then placed in their homecage. Half an hour later, their stillness was monitored for 30 min. Every four days, this procedure was repeated for a total of three times. The percentage of stillness was measured and averaged between the three repetitions for each mouse. A Mann-Whitney test yielded a significant difference between groups (p = 0.0333). Error bars represent the mean and the standard errors of the mean.

**Fig. S5.**
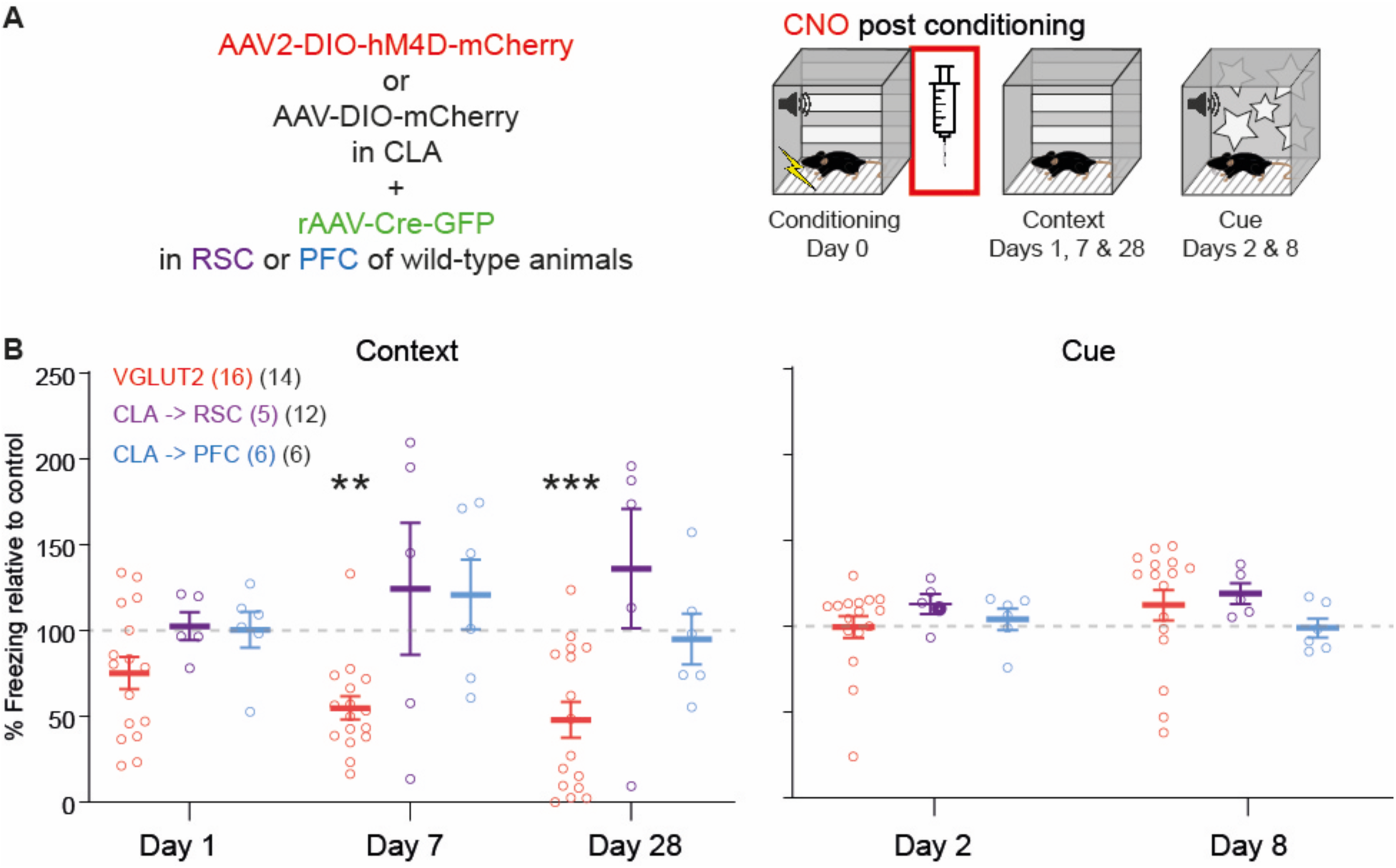
The inhibition of specific claustral projections does not impair the consolidation of a fear memory. (**A**) Using a retrograde transporting viral technique in wild type animals, claustral projections to the prefrontal (PFC) and retrosplenial (RSC) were inhibited (hM4D) during the consolidation phase after fear conditioning. CNO (2mg/kg, i.p) was injected right after conditioning to both hm4D and mCherry treated mice, then every 2.0 hours for a total of four injections. (**B**) Percentage of time spent freezing during re-exposures to the CC (left panels) and the CS (right panels) compared to control animals. For each experimental animal, its percentage of time spent freezing was divided by the average freezing time of all control animals (cage mates) tested the same day. However, comparisons were performed on the percentage freezing time between experimental groups and their respective controls. On days 1 (2), 7 (8) and 28 after conditioning, the freezing response to the conditioned context (stimulus) was monitored and compared to the freezing responses of mCherry treated mice. No difference or trends was observed between groups. For each group and each type of retrieval (context or cue), a two-way repeated measures ANOVA (treatment x days) was performed. CLA->PFC group during context retrievals yielded an effect of days (F(2) = 14.52, p = 0.0001) but no effect of treatment (F(1) = 0.7196, p = 0.4161) nor an interaction between treatment and days (F(2) = 0.1246, p = 0.8835). For the CLA->PFC group during cue retrievals, we found no effect of days (F(1) = 0.3078, p = 0.5912), of treatment (F(1) = 0.04977, p = 0.828) and no interaction (F(1) = 0.5806, p = 0.4637). For the CLA->RSP group during context retrieval, an ANOVA yielded an effect of days (F(2) = 9.527, p = 0.0006) but not of treatment (F(1) = 1.123, p = 0.306) nor an interaction (F(2) = 1.12, p = 0.3394). For the CLA->RSC group during cue retrievals, we found no effect of days (F(1) = 0.005958, p = 0.9395), nor treatment (F(1) = 2.069, p = 0.1709) and no interaction (F(1) = 0.1179, p = 0.736).

**Fig. S6.**
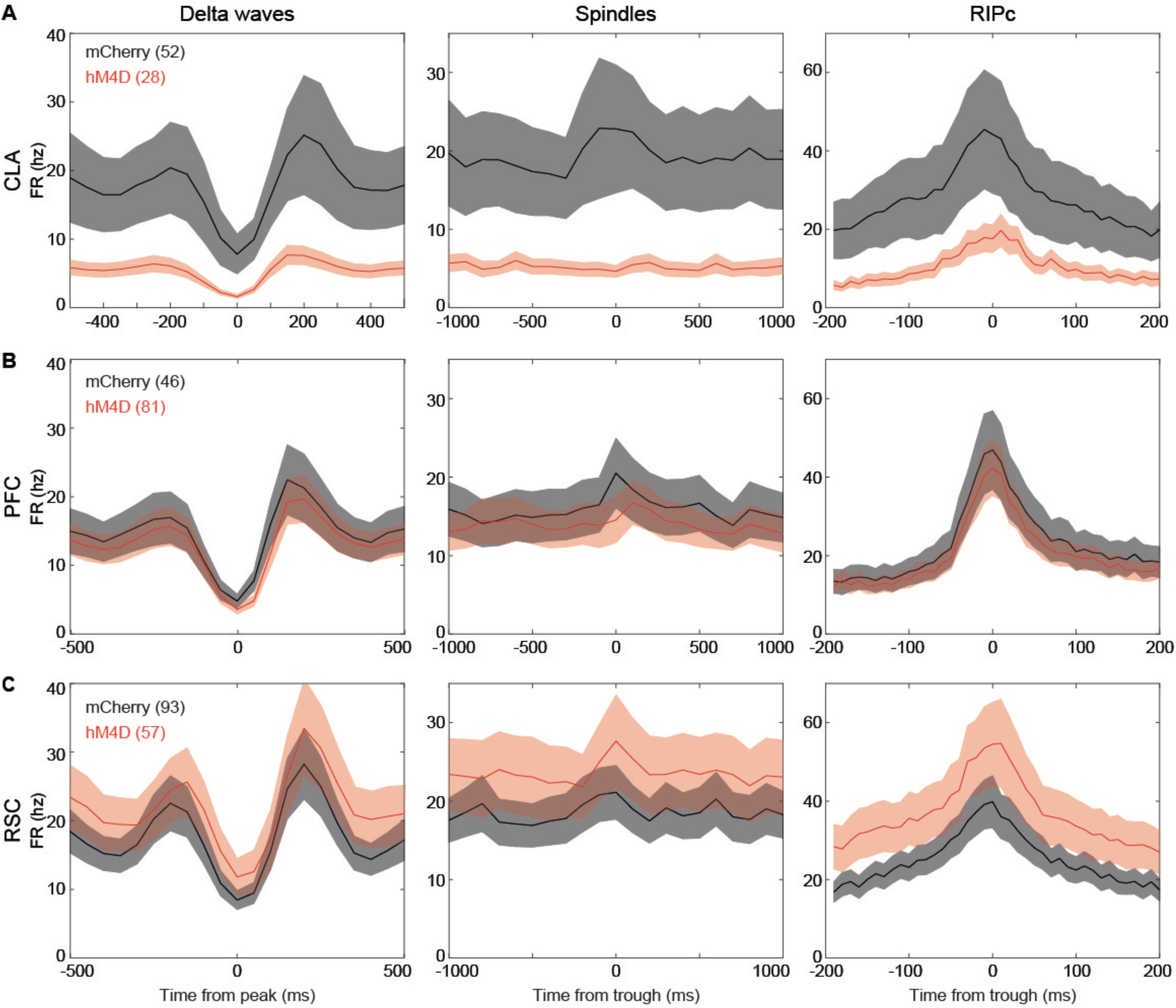
Neuronal spiking during delta waves, spindles and ripple-like events in the claustro-insular region, in the prefrontal cortex and in the retrosplenial cortex. Population peri-event time histograms during delta-waves (A), ripple-like events (RIPc; B) and spindles (C) for the claustro-insular region (CLA; left panels), the prefrontal cortex (PFC; middle panels) and the retrosplenial cortex (RSC; right panels). Lines and shaded areas correspond to the mean plus and minus the standard error of the mean.

## Movies

Movie S1. Behavioral responses to the conditioned context. Thirty-five minutes after an injection of CNO, a control wild-type mouse (mCherry) is exposed to the conditioned context on day 28 after conditioning.

Movie S2. Inhibition of claustro-retrosplenial projections and behavioral responses to the conditioned context. Thirty-five minutes after an injection of CNO, a wild-type mouse (hM4D) is exposed to the conditioned context on day 28 after conditioning.

